# Alignment-Free Approaches for Predicting Novel Nuclear Mitochondrial Segments (NUMTs) in the Human Genome

**DOI:** 10.1101/239053

**Authors:** Wentian Li, Jerome Freudenberg, Jan Freudenberg

**Affiliations:** The Robert S. Boas Center for Genomics and Human Genetics, The Feinstein Institute for Medical Research, Northwell Health, Manhasset, NY, USA; Regeneron Genetics Center, Regeneron Pharmaceuticals, Inc, Tarrytown, NY, USA

**Keywords:** alignment-free, k-mer, mitochondria, NUMT, Jensen-Shannon divergence, Manhattan plot

## Abstract

The nuclear human genome harbors sequences of mitochondrial origin, indicating an ancestral transfer of DNA from the mitogenome. Several Nuclear Mitochondrial Segments (NUMTs) have been detected by alignment-based sequence similarity search, as implemented in the Basic Local Alignment Search Tool (BLAST). Identifying NUMTs is important for the comprehensive annotation and understanding of the human genome. Here we explore the possibility of detecting NUMTs in the human genome by alignment-free sequence similarity search, such as k-mers (k-tuples, k-grams, oligos of length k) distributions. We find that when k=6 or larger, the k-mer approach and BLAST search produce almost identical results, e.g., detect the same set of NUMTs longer than 3kb. However, when k=5 or k=4, certain signals are only detected by the alignment-free approach, and these may indicate yet unrecognized, and potentially more ancestral NUMTs. We introduce a “Manhattan plot” style representation of NUMT predictions across the genome, which are calculated based on the reciprocal of the Jensen-Shannon divergence between the nuclear and mitochondrial k-mer frequencies. The further inspection of the k-mer-based NUMT predictions however shows that most of them contain long-terminal-repeat (LTR) annotations, whereas BLAST-based NUMT predictions do not. Thus, similarity of the mitogenome to LTR sequences is recognized, which we validate by finding the mitochondrial k-mer distribution closer to those for transposable sequences and specifically, close to some types of LTR.

## Introduction

Mitochondrial DNA or the mitochondrial genome (mitogenome) is the genetic material of mitochondria. The human mitogenome is double stranded circular DNA with roughly 16,600 base pairs, containing 13 genes coding for proteins, and 24 for RNAs (2 rRNAs and 22 tRNAs). Each cell contains hundreds to thousands copies of the mitogenome (Bendich, 1987) depending on the function of the cell (Veltri et al., 1990). The energy-hungry cells in muscle and heart tend to have more copies of mitogenome than other cells (Torres, 2018). The replication of mitogenomes is independent from that of the nuclear genome (Bogenhagen and Clayton, 1977; Holt and Reyes, 2012; Clay Montier et al., 2009). Cancer cells may have less copies of mitogenome per cell, to be consistent with a lower level of cellular respiration and high level of fermentation (Warburg effect) (Reznik et al., 2016). Mitochondria are known to be maternally inherited, but paternal or a mixture of both maternal and paternal inheritance have also be observed (Schwartz and Vissing, 2002; Luo et al., 2018). Both mitochondria organelle and the mitochondrial DNA play an important role in certain human diseases (Wallace, 2018). Despite the complexity of its copy number as well as its distribution across cell types, its dynamics, its inheritance pattern, and its variability (Parsons et al., 1997; Van Der Walt et al., 2003), the mitogenome can be represented through a single reference sequence, which we use here for our analyses.

On this background, the endosymbiosis hypothesis states that mitochondria have a bacterial ancestry and co-evolve with their eukaryotic host cells, benefitting both entities by this co-existence (Margulis, 1970). Such a co-existence implies a molecular interaction between the two units. Early research had accordingly found hybridization of nuclear and mitochondrial DNA fragments in mouse (Du Buy and Riley, 1967). Driven by the curiosity that ATPase genes are encoded in both mitochondrial DNA in *S. cerevisiae* and in nuclear DNA in a related species, *N. crassa*, led to the discovery that *N. crassa* has both copies in mitochondrial and the nuclear genome (Van Den Boogaart et al., 1982). A series of early reports confirmed widespread homology between DNA segments from the mitochondrial and nuclear genome (Farrelly and Butow, 1983; Gellissen et al., 1983; Wright et al., 1983; Jacob et al., 1983; Kemble et al., 1983; Hadler et al., 1983; Tsuzuki et al., 1983). This widespread homology was explained by a process of integration of mitochondrial DNA into the nuclear genome. Following (Lopez et al., 1994), we here use the term “nuclear mitochondrial DNA segments/sequence” or NUMT, to refer these pieces of mitochondrial (MT) DNA or, mitogenome), which were inserted into the nuclear DNA, without requiring any amplification event after the nuclear insertion (Lopez et al., 1994).

The presence of NUMTs in the human genome (Herrnstadt et al., 1999; Mourier et al., 2001; Tourmen et al., 2002; Woischnik and Moraes, 2002; Ricchetti et al., 2004; Hazkani-Covo et al., 2010) is also of interest because it can be a type of repetitive sequence, which can contribute to the redundancy problem in sequencing, mapping, and genotyping (e.g., (Weber and Myers, 1997; Green, 1997; Derrien et al., 2012; Lee and Schatz, 2012; Li et al., 2014; Li and Fredenberg, 2014)). Variants within NUMTs are easily mistaken as MT variants (for a mitogenome variant database, see, e.g. (Preste et al., 2018) in genetic disease mapping studies (Wallace et al., 1997; Parr et al., 2006; Yao et al., 2008). Carefully designed protocols have been proposed to remove the “contamination” from NUMT to mitogenome variants calls (Ring et al., 2018).

NUMTs may also be viewed analogously to structural changes of the nuclear genome called segmental duplication (ancestral), or copy number variation (CNV) (germline, recent and within species), or copy number alteration (CNA) (somatic, such as cancer cells): which are all dynamic processes with different time scales. Accordingly, more recent MT insertions can either serve as genetic marker (Thomas et al., 1996; Caro et al., 2010; Lang et al., 2012; Dayama et al., 2014), or be functional (Willett-Brozick et al., 2001; Turner et al., 2003; Goldin et al., 2004; Schon et al., 2012). Somatic MT insertions in cancer cells can offer novel insight on genomic instability in cancer (Srinivasainagendra et al., 2017; Singh et al., 2017). The latter two types might be called de novo NUMT and somatic NUMT, respectively. Furthermore, NUMTs also provide an opportunity to study the evolutionary history of genomes (e.g., (Zischler et al., 1995; Perna and Kocher, 1996; Bensasson et al., 2001; Mishmar et al., 2004; Gunbin et al., 2017)).

Several attempts were made to catalog all human NUMTs. After the initial human reference genome became available, 296 NUMTs were identified through a combination of alignment (BLAST with the default setting) and co-linearity of conserved blocks, with sizes between 106 bp and 14kb (Mourier et al., 2001). Comparisons of various attempts of compiling human NUMTs found 190 consistent entries (Lascaro et al., 2008). The current standard list of human NUMT is based on two studies with very similar number of hits: 755 NUMTs in (Ramos et al., 2011) and 766 NUMTs in (Simone et al., 2011), with an update to 585 NUMTs in (Calabrese et al., 2012).

This list of computationally predicted (*in silico*) NUMTs is based on sequence alignment, in particular using BLAST alignments (Altschul et al., 1990). The BLAST algorithm starts by finding exactly matching k-mers (l-tuples, n-grams) in two sequences, with k usually being small. Any matching k-mer is then extended by a local alignment to its maximum length. The aligned segments, with possible gaps and mismatches, are then returned as BLAST alignment hits. The biological assumption behind the design of this algorithm is that the divergence between two sequences is exclusively caused by small insertions, deletions, and point mutations. However, if the mutational dynamics of the sequences have been more complicated, such alignment-based approaches like BLAST may fail to detect significant similarity between two sequences (many mismatches due to greater evolutionary distance is another situation where alignment-based methods may fail).

To overcome these challenges, alignment-free methods have been proposed (Blaisdell, 1986; Vinga and Almeida, 2003; Son et al., 2014; Luczak et al., 2017; Ren et al., 2018), as these do not consider the ordering of the matching k-mers in two sequences to detect their similarity, and, possibly more remote homology. Another potential problematic situation for alignment-based methods are hot spots of insertion of short DNA segments, where the insertion occurs at different time and at random insertion points. In these instances, even if the ancestry of DNA segments were identical, the lack of ordering of the inserted pieces would prevent an alignment-based to detect the homology.

In this paper, we ask the question whether large BLAST-alignment-detected NUMTs can be equivalently detected by alignment-free methods, i.e., can be recognized by the similarity of k-mer/l-tuple distributions. Previous analyses have found some notable differences in the predicted amount of repetitive sequences in the human genome with alignment-based versus alignment-free methods, with repetitive sequences accounting for one-half with the former and accounting for two-third with the latter (Gu et al., 2008; De Koning et al., 2011). Therefore, we may expect the possibility of different results between alignment-free k-mer-based and alignment-based BLAST searches for NUMTs. Our paper is organized as follows: we first describe our k-mer analysis design; we then establish that k-mer distributions between MT and nuclear genome are indeed distinguishable; k-mer based NUMTs detection is then carried out for k=6,7, and for k=4,5; and the results are discussed in the context of our current knowledge about NUMTs as well as the broader biological context.

## Results

### Issues in designing a k-mer based NUMT detection method

We apply a moving window to scan the nuclear genome sequence for regions with a similar k-mer frequency distribution as the mitochondrial genome sequence. There are several decisions to be made when implementing a practical detection approach:

1. **Window size and step size:** The choice of window size is closely related to the size of MT genome. Here we chose a window size of 3kb, and moving step size of 1/8 of the window size, targeting NUMTs in the range of 3kb size. In order to test the robustness of our results, we also rerun calculations at peak locations with a moving step size of 1 base. The size of the human mitochondrial genome is 16,569 bp, and thus we expect to detect NUMTs that comprise roughly one fifth of its size.
2. **Measure of similarity between two k-mer frequency distributions:** This topic is extensively covered in (Vinga and Almeida, 2003). We use the Jensen-Shannon (JS) divergence (Burbea and Rao, 1982; Rao, 1982; Lin, 1991) based on an application of Jensen’s inequality for convex/concave functions. We denote {*m*_*i*_ = (*p*_*i*_ + *q*_*i*_)/2} for the high-dimensional mid-point between the two distributions, and JS divergence is defined as: 
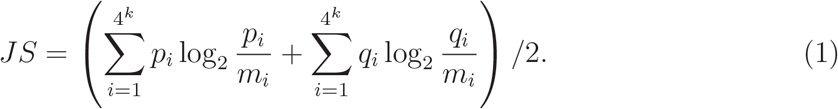 JS divergence is a symmetric version of the Kullback-Leibler distance (relative entropy). We previously used JS divergence to segment DNA sequences into relatively homogeneous halves (Bernaola-Galvan et al., 1996; Grosse et al., 2002; Li, 2001; Li et al., 2002) The alignment-free similarity is measured by the reciprocal of JS divergence: 1/JS. Other measures have been tried, such as the Euclidean distance, but JS divergence proved to be a choice that was more consistent with alignment-based approaches.
3. **The value(s) of k**: We are using the following k values: 3,4,5,6,7, based onn consistent findings from several procedures for choosing the value of k. The first is to find the k value that maximizes the number of more-than-once k-mer types (Sims et al., 2009). The idea behind this approach is that if a sequence is constructed with each k-mer type occurring only once (Fraenkel and Gillis, 1966) (a Hamiltonian cycle on the De Bruijn graph where each k-mer is represented by a node (e.g., (Pevzner et al., 2001))), no k-mer type could claim to represent the sequence. Only k-mer types which appear more than once have a chance to be part of the signature, and it is desirable to maximize the set of k-mer types. Fig.1(A) shows that for MT sequence, the number of k-mer types (with one or more counts) peaks at k=7. Another strategy for choosing the value for k is based on whether a k-mer type frequency is well predicted by all (k-1)-mer and (k-2)-mer frequencies (Sims et al., 2009). If the predicted and observed k-mer type frequencies are close (measured by the Kullback-Leibler (KL) divergence), there is not much new information gained by increasing the oligo length to k. Fig.1(B) shows that the KL divergence is highest around k=7-8. (and reaches zero at k=14). Yet another consideration for choosing k is provided by the formula: k=log_4_(*N*) + 1 (Price et al., 2005; Campagna et al., 2004; Gu et al., 2008), where *N* is the size of the region where k-mer frequency is counted, which can be the whole genome length, one chromosome length, or window length. This number is very similar to the formula k=log_4_(2*N*) = log_4_(*N*) + 0.5 when each k-mer type appears only once in the genome (Li et al., 2014). Based on these two formulas, for a 3kb window, the optimal choice of k is 6.28-6.78, and for 12,000 base MT sequence, the choice of k is 7.28-7.78. Even longer k-mers are considered in (Wang et al., 2016), serving different purposes (and not discussed here). Previous empirical attempts to choose an appropriate *k* value can be found in (e.g.) (Chor et al., 2009; Zuo et al., 2014; Jia et al., 2018) for different applications.
4. **Considering strand asymmetry instead of combining direct and reverse-complement k-mers**: Strand symmetry (or Chargaff’s second parity rule) (Elson and Chargaff, 1952; Li, 1997; Forsdyke, 2016) is true for k-mers (Prabhu, 1993) for most long genomic sequences. If the strand symmetry holds, a k-mer (e.g. AACGT) has almost identical frequency as its reverse complement (e.g. ACGTT), then these two k-mers might be grouped into one unit. However, mitochondrial sequence is short, and many organellar sequences do violate strand symmetry (Nikolaou and Almirantis, 2006). The strand asymmetry in mitochondria is the basis for the distinction between heavy and light strands (e.g. (Reich and Luck, 1966)). Consequently, we use the frequency of all 4^*k*^ k-mer types, not 4^*k*^ /2, in characterizing the MT sequence.
5. **Not normalizing the k-mer frequencies by (k-1)-mer frequencies:** One version of using k-mers to characterize a genomic region or a genome is to compare the observed k-mer frequencies with the expected frequencies as predicted from the (k-1)-mer frequencies, and use the ratio of the two as a measure. This approach was called “genomic signature” in (Karlin et al., 1997; Campbell et al., 1999). Our experiment shows that un-normalized k-mer frequencies perform better than the normalized ones. We note however that other versions of normalization such as subtracting the background/expected frequencies (Wan et al., 2010), instead of ratio, are not tested.
6. **Threshold for calling NUMTs at similarity peaks of k-mer distributions:** When 1/JS is high enough (or JS low enough), we make a NUMT prediction call. The threshold of the peak call was based on previously identified NUMTs that are longer than 3kb (same as the window size). The 1/JS values of all these large NUMT loci are sorted, and the second lowest 1/JS value is used as the threshold. Since there are 38 NUMTs in the most recent collection (Calabrese et al., 2012) that are larger than 3kb, missing 2 peak calls correspond to a false negative rate of 5%.

**Figure 1:**
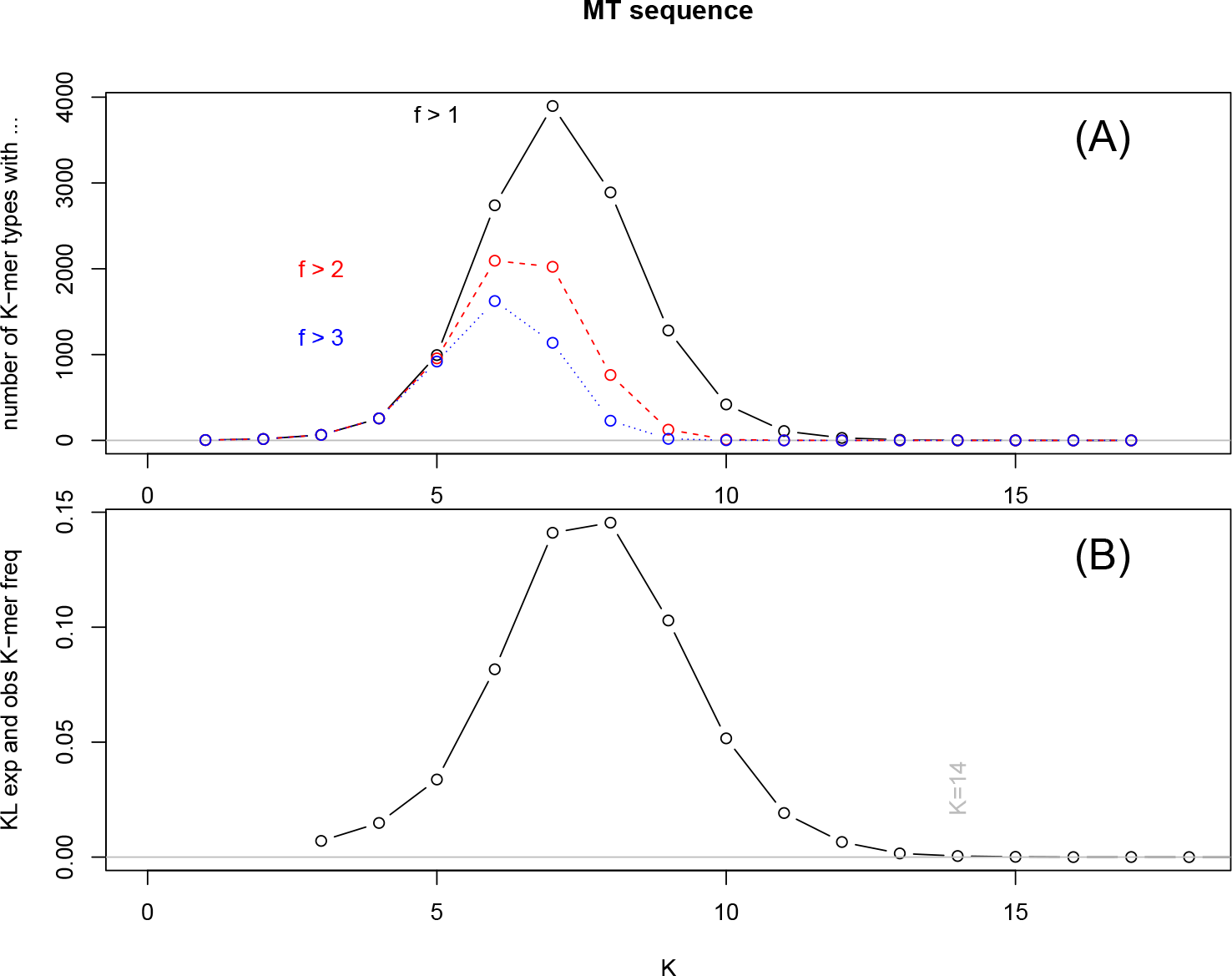
(A) The number of k-mer types in human MT sequence that appear at least once, as a function of k. The red and red points denote the number of k-mer types that appear at least two and three times in the MT sequence. (B) The Kullback-Leibler divergence between the observed k-mer frequencies and those predicted by (k-1) and (k-2)-mer frequencies (Sims et al., 2009).

Note that we did not use other public domain, specially-designed, computationally-efficient k-mer count programs, because the small k values chosen in this paper allowed for less optimal implementations. Many space/time efficient programs have been developed for dealing with much larger k-values (e.g. k > 20), and one can refer to these publications: (Kurtz et al., 2008; Marcais and Kingsford, 2011; Melsted and Pritchard, 2011; Rizk et al., 2013; Deorowicz et al., 2013; Zhang et al., 2014; Roy et al., 2014; Patro et al., 2014; Audano and Vannberg, 2014; Melsted and Halldorsson, 2014; Deorowicz et al., 2015; Mamun et al., 2016; Sivadasan et al., 2016; Bray et al., 2016; Pandey et al., 2017; Marchet et al., 2017; Ebert et al., 2017; Kokot et al., 2017; Patro et al., 2017).

### Nuclear DNA and mitochondrial DNA have overall different k-mer distributions

Different k-mer distributions of the nuclear genome sequence and the MT sequence are already expected based on the difference in GC-content: the GC-content of human MT sequence is 55.6%, as compared to the GC-content in the nuclear genome of around 40% (Li, 2013). To further illustrate that the k-mer frequency of the nuclear and MT genome are different, we use a 3kb window to scan 5-mer frequencies in the first 10Mb region of human chromosome 1 (non-overlapping windows), as well as the first 15kb region of mitochondrial DNA.

Fig.2(A) shows points representing the windows in the first two dimensions of multidimensional scaling (MDS). The MT is not the most distinct sequence in comparison to the genomic sequence: a low-complexity region (chr1:2.652Mb-2.772Mb, hg38) is the most obvious outlier. Windows from MT genome are near the outskirt of the main band. When windows with mostly (more than 99%) unique (non-repetitive) sequences and those with mostly (more than 99%) repetitive sequences are marked (red and green, respectively), MT windows are closer to repetitive sequences. Fig.2(B) shows the second and third dimension of MDS, and again MT windows are on the border of the main band. Also from Fig.2(A,B) note that windows belong to MT are tightly clustered, indicating that in the context of nuclear genome, MT sequence is relatively homogeneous.

**Figure 2:**
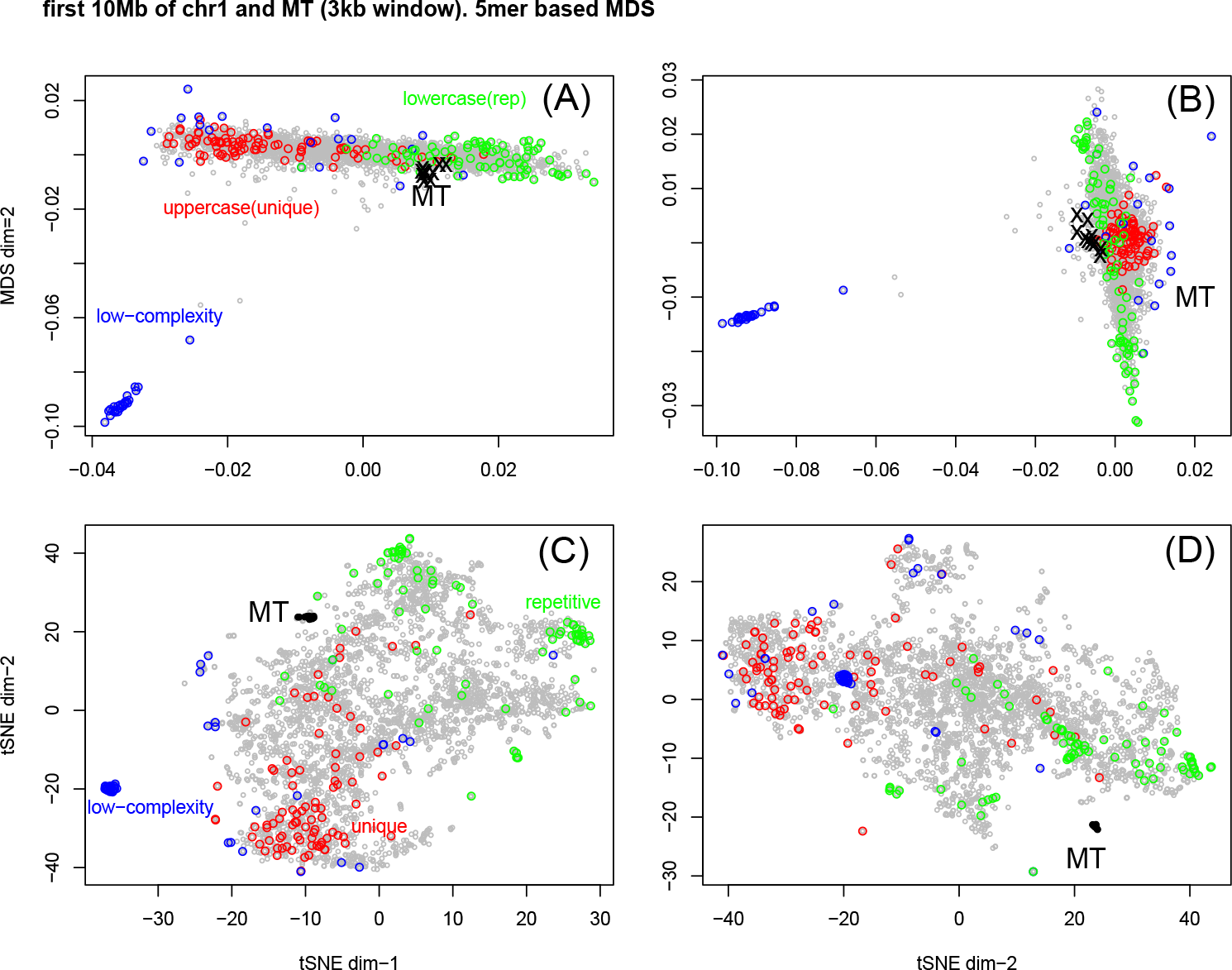
Multi-dimensional scaling (MDS) for (A) the 1st and 2nd dimension; and (B) 2nd and 3rd dimension. Each point represents a 5-mer frequency distribution of a 3kb window (non-overlapping window in human chromosome 1, overlapping in human mitochondrial DNA). Low complexity windows (E < 8.1) are marked in red. Windows with high proportion of non-repetitive (unique, not filtered by RepeatMasker, uppercase) bases are marked in red, and those with high proportion of repetitive sequences (lowercase) are marked in green. The MT windows are marked in black. (C) and (D) are equivalent tSNE plot of the 1st vs 2nd dimension, and 2nd vs 3rd dimension.

To seed whether the MDS dimensions (1,2,3) represent simple features of the sequence, we calculate the following parameters: percentage of unique (non-repetitive) sequence, GC-content, Jensen-Shannon divergence between the k-mer frequency and its reverse-complements. The latter is a measure of the strand asymmetry at the k-mer level: zero means strand symmetry, and large value denote violation of strand symmetry. Table 1 shows that MDS dimension-1 closely follows the GC-content (Spearman correlation coefficient (cc) = −0.98, p-value < 2 × 10^−16^). The negative sign doesn’t have any particular meaning, as the direction of the MDS axis is arbitrary. MDS1 is also highly correlated with proportion of unique sequences (Spearman cc = −0.66, p-value < 2 × 10^−16^) MDS1 is not only correlated with these two quantities again (though with lesser strength), but also weakly correlated with the strand asymmetry (Spearman cc= −0.1, p-value = 2 × 10^−8^). MDS3 is weakly correlated with GC-content (Spearman cc= 0.055, p-value = 0.002).

**Table 1:** Spearman’s correlation coefficient between simple summary statistics of sequence windows (proportion of unique sequences, GC content, Jensen-Shannon divergence between 5-mer frequencies and those of its reverse complements) and the first three dimensions from the multi dimensional scaling (MDS) analysis across all windows.

|  | MDS1 | MDS2 | MDS3 |
| --- | --- | --- | --- |
| % unique | -0.66 (pv=0) | 0.32 (pv=0) | 0.02 (pv=0.24) |
| GC | -0.98 (pv=0) | 0.58 (pv=0) | 0.055 (pv=0.002) |
| strand asym | 0.04 (pv=0.02) | -0.098 (pv=2E-8) | 0.017 (pv=0.3) |

The classic/metric MDS is a linear projection from high to low dimensional space. A newer dimension reduction and plotting technique, tSNE (t-distributed stochastic neighbor embedding), is a nonlinear projection (Van Der Maaten and Hinton, 2008). Previously, we applied tSNE to human genetic data (Li et al., 2017), and found it quite powerful in displaying both continental and sub-continental distributions of genetic variation. Fig.2(C)(D) shows the tSNE for 1st vs 2nd dimension, and 2nd vs 3rd dimension. Unlike MDS, tSNE is able to display the substructure within the main band, even with the presence of the outliers (the low-complexity sequence). It is clear that MT points for a cluster being separated from the rest of the genomics sequences in chromosome 1. A close inspection of the plots shows three chr1 windows within the MT cluster, which are predicted as being NUMTs.

### Manhattan plot of alignment-free peaks for NUMTs at k=6 and k=7

A common technique in genome-wide association studies (GWAS) of human genetically complex traits are the so called Manhattan plots, which plot genetic variant association signals (often measured by the −log(p-value)) over their genomic position across all chromosomes (e.g. (Wellcome Trust Case Control Consortium, 2007)). Here we introduce this idea for plotting the alignment-free signal of mitochondrial DNA insertion across the nuclear genome. The signal strength plotted on the *y*-axis is the reciprocal of JS divergence, 1/JS, between the two k-mer distributions, with the target being the overlapping window of size 3kb along the chromosome, and with the query being the mitochondrial sequence (and its reverse complement).

Fig.3 shows the Manhattan plot for k=6 (only windows with high enough signal are included). First of all, regions with the highest peak heights are easily identified (chr1, chr5, chr17), indicating a high level of similarity with the MT sequence. Second, all peaks (determined by 1/JS > 3.346) can be explained by the known NUMTs in (Calabrese et al., 2012), with sizes larger than 3kb (pink), 2kb (blue), or 1kb (green). In other words, at k=6, alignment-free approach does not reveal any new potential NUMTs which were not already identified by the alignment-based BLAST method.

**Figure 3:**
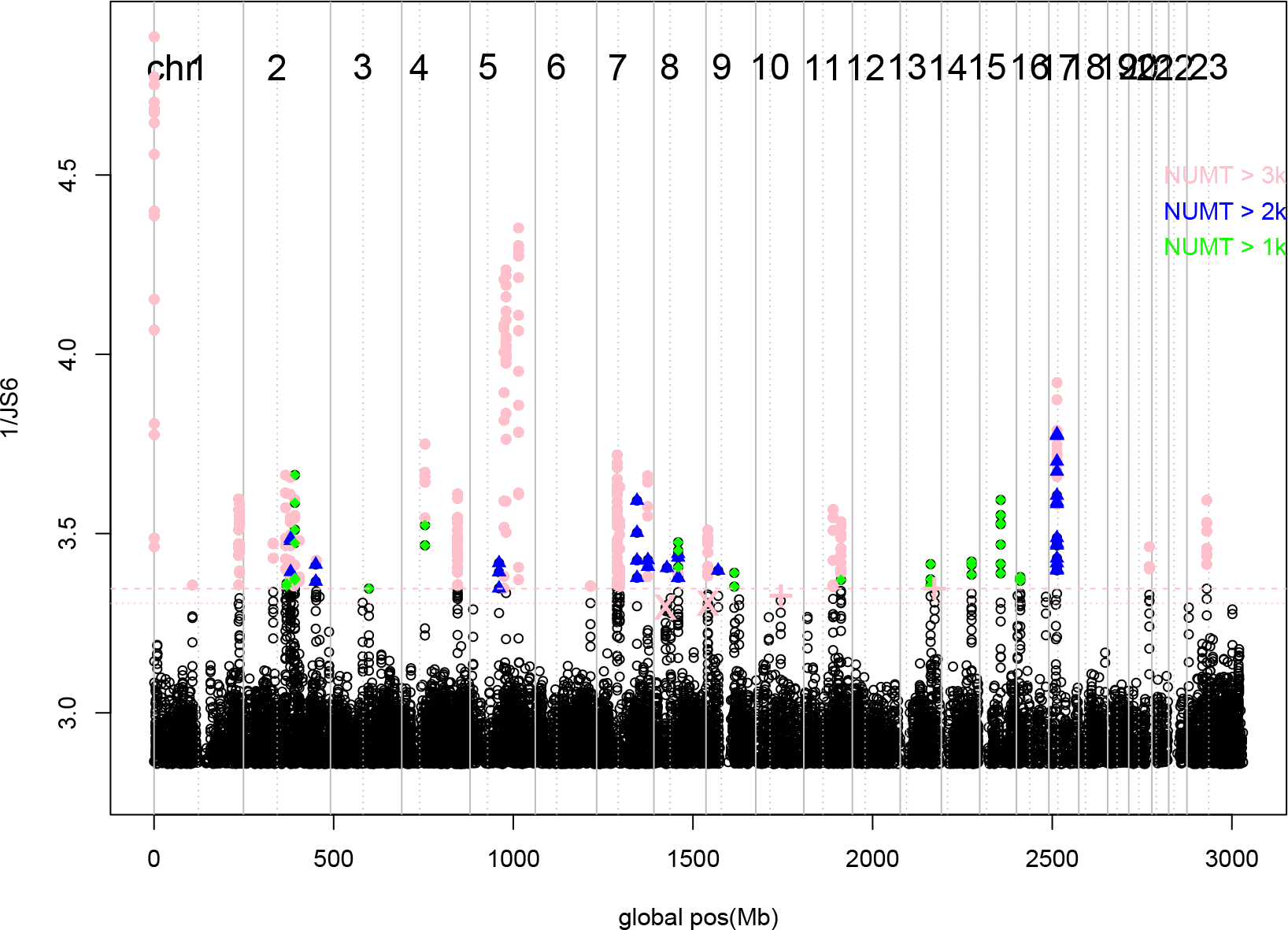
Manhattan plot of alignment-free NUMT detection signals, as obtained by 1/JS for k=6, and plotted as a function of the global genomic location (Mb). Each point represents a 3kb window, which moves by 1/8 of the window size (375 bases). Windows with 1/JS being less than 1/0.35=2.857 are not plotted. Chromosome start/end positions are marked by vertical solid lines, and centromeres marked by vertical dashed lines. The horizontal lines represent the threshold for peak calling: the threshold value for calling a NUMT prediction is chosen to be the second lowest 1/JS values among 38 previously known NUMTs (with size larger than 3kb). The difference between the two horizonal lines is that one was obtained from moving windows 375 bases (for 1/JS=3.346), and another from moving windows by 1 base (for 1/JS=3.305). All observed peaks above these lines can be explained by the known NUMTs as listed in (Calabrese et al., 2012): those with sizes larger than 3kb are colored in pink, those with size between 2kb and 3kb in red, and those between 1kb and 2kb in green.

We note that some peaks are very close to either telomere (chr1) or centromere (chr17). It has been observed that somatic large-scale structural changes in cancer cells can be grouped into either the telomere/centromere-bounded or not-telomere/centromere-bounded category (Zack et al., 2013). The two categories display different sizes of structural variants, and may be caused by different mechanisms. In Fig.3, two out of the four highest peaks are close to telomere or centromere.

Our observations that for k=6 alignment-free peaks are fully explained by alignment-based NUMTs at 3kb or larger, as well as some NUMTs with 1kb-3kb sizes, shows the robustness of our choice for NUMT calling. In the initial search, peaks are called based on the second lowest 1/JS value for the 38 NUMTs larger than 3k, when 3kb window moves 375 bases at the time (that 1/JS is 3.305). We do not come to different conclusions, when we move up the threshold for peak calling at 1/JS > 3.346, or when the window moving step size is reduced from 375bp to 1 bp. To test the robustness of our parameter choices, we also lowered the threshold for peak calling by the lowest (instead of the second lowest) 1/JS value for the 38 NUMTs larger than 3kb (1/JS > 3.2945): again, all peaks are explained by known NUMTs 1kb or larger.

Similar conclusion are also reached for k=7 (plot not shown). If we use 1/JS > 7.7336 as the peak calling criterion (which is the second lowest 1/JS value for +3kb NUMTs with moving step size of 1 base), all peaks are also identified as by BLAST-based NUMTs of 1kb or larger. When the peak calling is less stringent, at 1/JS > 1.711 (lowest 1/JS value for +3kb NUMTs with moving step of 1 base), 1.6845 (second lowest 1/JS value with moving step of 375 bases), 1.659 (lowest 1/JS value with moving step of 375 bases), we either observe the same conclusion or only a few isolated probably false-positives.

### New alignment-free peaks with k=5 and k=4

New alignment-free similarity peaks can be found when k is reduced to 5 or below. This is shown in Fig.4 which has 9 loci with peaks (1/JS > 8.666, which is the second lowest peaks for known NUMTs larger than 3kb, with moving step size of 1 bp) that do not have an underlying known NUMTs of 1kb or longer. Two of the loci (chr3:89.587Mb, chrX:126.471Mb) contain known NUMTs of length shorter than 1kb. The remaining 7 loci are listed in Table 2.

**Figure 4:**
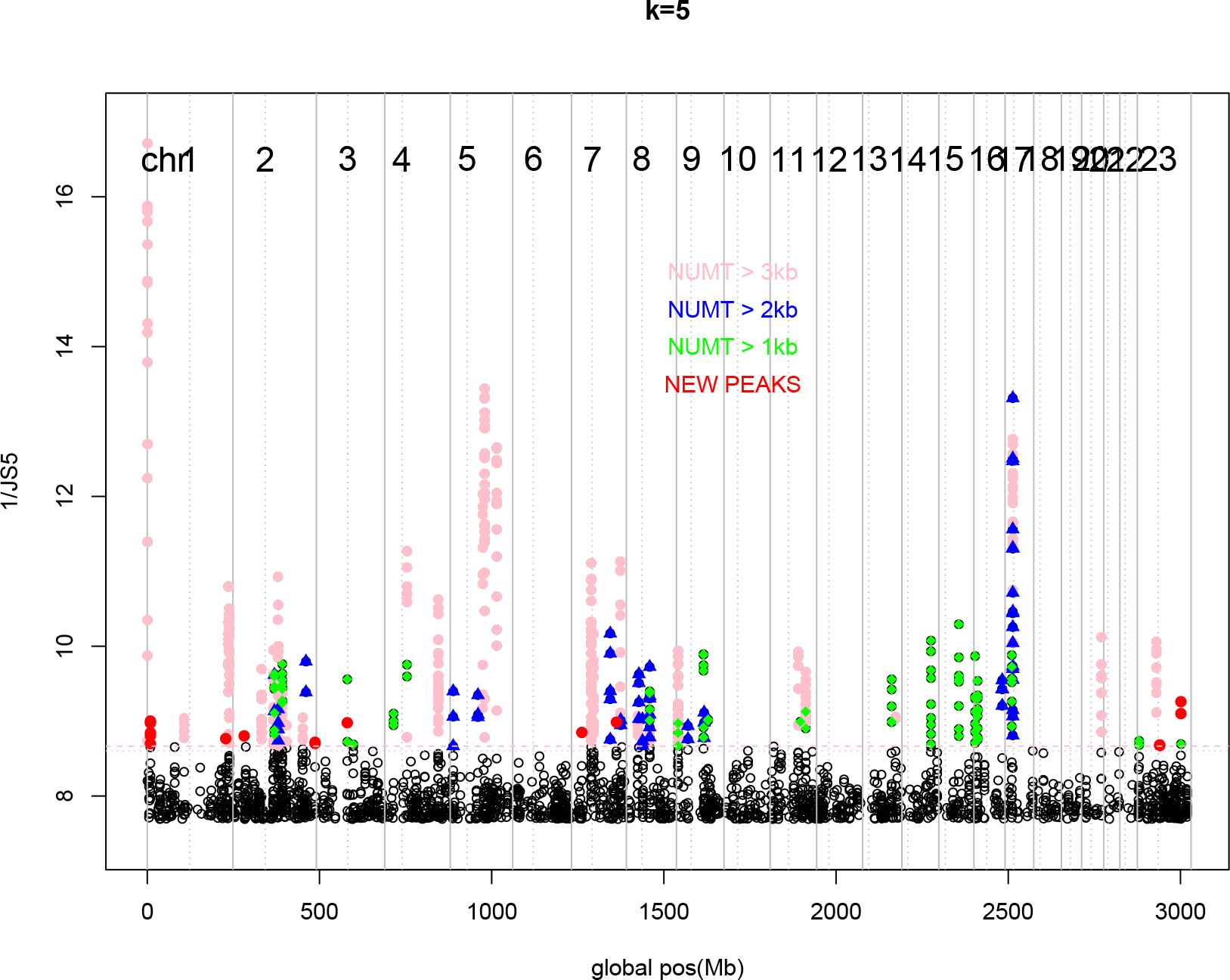
Manhattan plot of alignment-free NUMT detection signals, as obtained by 1/JS for k=5, again plotted as a function of the global genomic location (Mb). Each point again represents a 3kb window, which moves by 1/8 of the window size (375 bases). Windows with 1/JS less than 1/0.14 =7.143 are not plotted. The horizontal lines represent the threshold for calling NUMTs from peak signals: the second lowest 1/JS values among 38 known NUMTs with size larger than 3kb, 1/JS=8.604 for step size 375 and 1/JS=8.666 for step size 1. Peaks which are explained by known NUMTs listed in (Calabrese et al., 2012) of size larger than 3kb are colored in pink, those with size between 2kb and 3kb in red, and those between 1kb and 2kb in green. New NUMT prediction peaks are marked in red.

**Table 2:** List of NUMT predictions without a corresponding BLAST-based NUMT as called from similarity peaks for k=5 and 4. Column headers. k: the k value at which the peaks are detected; chr: chromosome; pos: chromosome position in Mb, hg38; 1/JS: similarity of k-mer distributions as measured by the reciprocal of Jensen-Shannon entropy; nw: number of 3kb windows that pass the threshold; annotation: other information obtained from the UCSC Genome Browser.

| k | chr | pos(Mb,hg38) | 1/JS | nw | annotation |
| --- | --- | --- | --- | --- | --- |
| 5,4 | 1 | 9.090375-9.094875 | 9.001, 24.938 | 5 | LTR |
| 5,4 | 1 | 227.635875-227.638875 | 8.767, 23.124 | 1 | LTR, gene ZNF678 |
| 5,4 | 2 | 32.357625-32.360625 | 8.803, 25.392 | 1 | gene BIRC6, DNase cluster |
| 5,4 | 2 | 238.206375-238.209750 | 8.718, 23.702 | 2 | LTR |
| 5 | 7 | 29.789625-29.792625 | 8.849 | 1 | LTR |
| 5,4 | 7 | 130.314375-130.317375 | 8.986, 26.431 | 1 | LTR, SINE, gene CPA4 |
| 5 | X | 64.846875-64.850250 | 8.682 | 2 | LTR |
| 4 | 3 | 141.696375-141.702250 | 24.198 | 6 | LTR |
| 4 | 4 | 145.835625-145.838625 | 23.346 | 1 | LTR |
| 4 | 6 | 17.319000-17.323125 | 23.942 | 2 | (LTR) |
| 4 | 8 | 47.988750-47.992500 | 23.869 | 3 | LTR |
| 4 | 11 | 122.828625-122.831625 | 23.214 | 1 | LTR, (LINE) |
| 4 | 13 | 54.939375-54.942375 | 23.938 | 1 | LTR |
| 4 | 13 | 83.908125-83.911125 | 23.268 | 1 | LTR |
| 4 | 14 | 102.244125-102.248625 | 26.094 | 5 | LTR, gene MOK |
| 4 | 16 | 30.541875-30.545250 | 24.845 | 2 | LTR, gene AC002310.13 |
| 4 | 17 | 80.401500-80.404500 | 23.731 | 1 | LTR, gene LOC100294362/CTD-2047H16.4 |
| 4 | 18 | 28.209375-28.212375 | 23.205 | 1 | LTR |
| 4 | X | 47.286375-47.289375 | 23.002 | 1 | LTR |

With new peak near chr1:9.093Mb being the highest and widest we plot the 1/JS in the region separately in Fig.5(A). The 3kb window size is marked for each point as a reminder of the length scale. If the peak height is about 9, and the genome-wide average is 5, the half-height points can be used to measure the peak width. This leads to the peak region from 9087.75kb to 9096.75kb, or, of width of 9kb. Other peaks listed in Table 2 have narrower widths, as well as lower heights.

**Figure 5:**
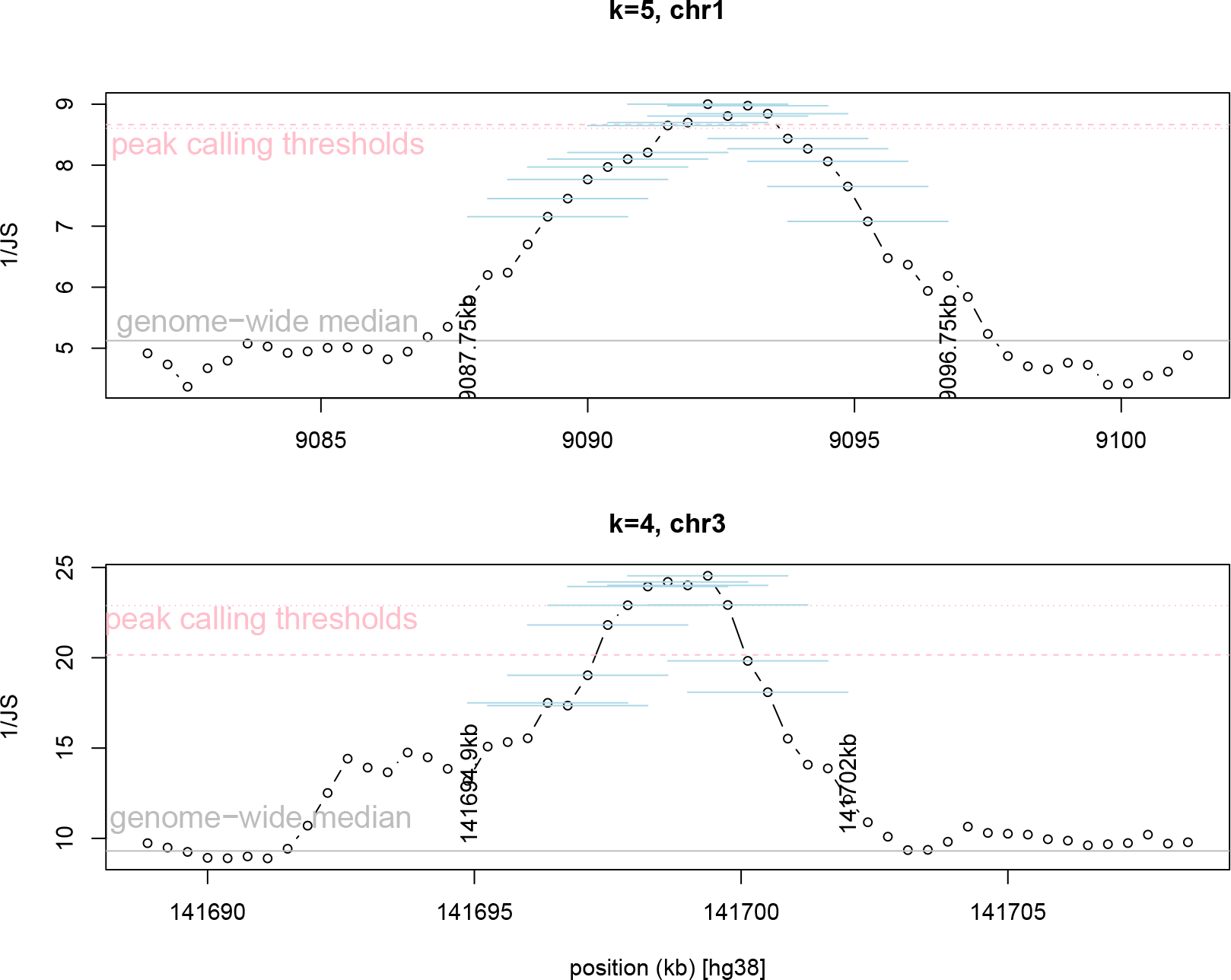
(A) Distribution of 1/JS for k=5 over a signal peak region at chromosome 1. The peak calling threshold is marked by pink, and genome-wide median of 1/JS is marked by grey. The peak width is estimated from the left end of the 3kb windows reaching 1/JS=7, to the right end of the 3kb window falling off from 1/JS=7, which is from 9087.75kb to 9096.75kb, or 9kb. (B) Distribution of 1/JS for k=4 over a signal peak region at chromosome 3. The width of the peak is estimated to be 7.1kb from 141694.9kb to 141702kb.

At k=4, the relative peak heights of +3kb NUMTs span a much bigger range. If we still use the second lowest peak value of the 38 +3kb NUMTs, many new loci will pass the peak call. To be more conservative, we use the 4th lowest value among the +3kb NUMTs, which is 1/JS = 22.89, as the threshold. With this criterion, 18 loci are called: 5 of them already appear for k=5; 1 with known underlying NUMTs of less than 1kb size. This leads to 12 new alignment-free signals at k=4 (see Table 2).

We arbitrarily picked one of the new alignment-free peaks for k=4 at chr3:141.699Mb to be examined in more detail (Fig.5(B)). If we consider the peak height as 1/JS=24, baseline as 1/JS=10, the half-height value is 1/JS=17. Using this half-height, the width of the peak is roughly 7.1kb (extending both ends of the 3kb window).

The alignment-free similarity signal for k=5/chr1 and k=4/chr3 are further analyzed by the pairwise BLAST program comparisons to the mitochondrial genome. As expected, there are no hit if megablast (highly similar) and discontiguous megablast (more dissimilar) options are used. The default options for seed word lengths are *k* = 28 and *k* = 18 respectively. However, blastn (with the default seed word length *k* = 11, expected threshold of 10, match, mismatch, gap, gap extension score to be 2, −3, −5 and 2) produce 9 (8) matches between the chr1 (chr3) peak and MT sequence.

In order to compare our alignment-free results with shorter *k*–mers with those by alignment-based methods, we re-run the BLAST program (online and command-line version with slightly different default settings, and turn off the RepeatMasker filtering) at several reduced seed word lengths. The run result is summarized in the Appendix. Both the online version and the command-line version have restrictions on choosing the *k* value: only three choices are given (7,11,15) for the online version, whereas *k* cannot be lower than 4 in the command line version. When *k* is reduced (while keeping the same E threshold), more alignments are detected, with only slight increase of the maximum alignment length, as well as a small increase of the total scores (sum of individual alignment scores). This indicates that the seed word length is not a crucial factor in using BLAST to detect a long alignment.

On the other hand, if the threshold of BLAST alignment is relaxed (to e.g., *E* = 20), there is a large increase in both the number of alignments and the total alignment score (see Appendix). The LAST program also confirms this observation. LAST does not require a seed word length because it starts from the seed of one base. Therefore, the main way to relax the alignment criterion is to use a smaller *D* parameter value. Indeed, smaller *D*s lead to both more alignments and longer alignment lengths (see Appendix).

### Annotation tracks of the new alignment-free peaks

We used the UCSC genome browser annotation to examine all +3kb BLAST-based NUMTs and new alignment-free peak regions listed in Table 2. Both the BLAST-based large NUMTs and new alignment-free peak regions overlap with genes: RP5-857K21.4 (chr1, HSA NumtS 001), MTRNR2L11 (chr1, HSA NumtS 043), ARHGAP15 (chr2,HSA NumtS 090), JAK2 (chr9, HSA NumtS 329), DNAJC3 (chr13, HSA NumtS 470), AQP9 (chr15, HSA NumtS 490), RAE1 (chr20, HSA NumtS 547), SMARCA2 (chrX, HSA NumtS 565), and ZNF678 (chr1), BIRC6 (chr2), CPA4 (chr7), MOK (chr14), AC002310.13 (chr16), LOC100294362/CTD-2047H16.4 (chr17) (Table 2). There is no indication of a difference in the tendency to co-localize with genes in the two groups.

Multiple large BLAST-based NUMTs overlap with DNase hypersensitive site clusters, in-cluding HSA NumtS 001, 042, 043 (on chr1), 219, 222, 228 (on chr5), 508 (on chr17). For our alignment-free peak regions, only one (on chr2) overlaps with DNase hypersensitive site clusters. There appear to be isolated appearance of SINE and LINE, but these are not prominent in these regions.

The biggest annotation difference between the BLAST-based and alignment-free peaks is the co-localization with Long Terminal Repeat (LTR), also called endogenous retrovirus (ERV) (Thompson et al., 2016). All alignment-free peak regions in Table 2, with one exception, cover a LTR, in particular, in the ERV1 family (note that it should not be confused with the yeast gene ERV1). On the other hand, only one BLAST-based NUMTs (HSA NumtS 565, chrX) covers a LTR. The difference is statistically very significant (χ^2^-test *p*–value - 3×10^−11^) indicating that biological character of NUMT predictions as derived from the two types of signals might be different.

To evaluate the relation between MT sequence and transposable elements more directly, we compare the k-mer (k=5) distributions between mitogenome and all repetitive sequences from the Repbase database (Jurka, 2000; Kojima, 2018). To make sure the sequence length does not create a bias, we use not only the mitogenome itself, but also the five 5kb non-overlapping segments, and 16 1kb segments, to represent MT. Fig.6(A,B) shows the Multi-dimensional scaling (MDS), and tSNE, of low-dimensional projection of 5-mer distributions of all repetitive sequences and various MT sequences. Fig.6(A) shows that points representing MT are surrounded by points representing LTR. On the other hand, non-LTR (e.g. LINE, SINE including Alu), DNA transposons, satellites, with some exceptions, are more distant. tSNE does not preserve distance and its purpose is to preserve nearest neighbors. It is then not surprising that MT sequences are close to each other. However, the point representing the first 3kb window (and three 1kb window points) are closer to a few points representing LTR sequences and/or DNA transposons. These observations combined with that of Fig.2, that MT is closer to transposons than to unique sequences, are further supporting evidence for a MT-LTR connection.

**Figure 6:**
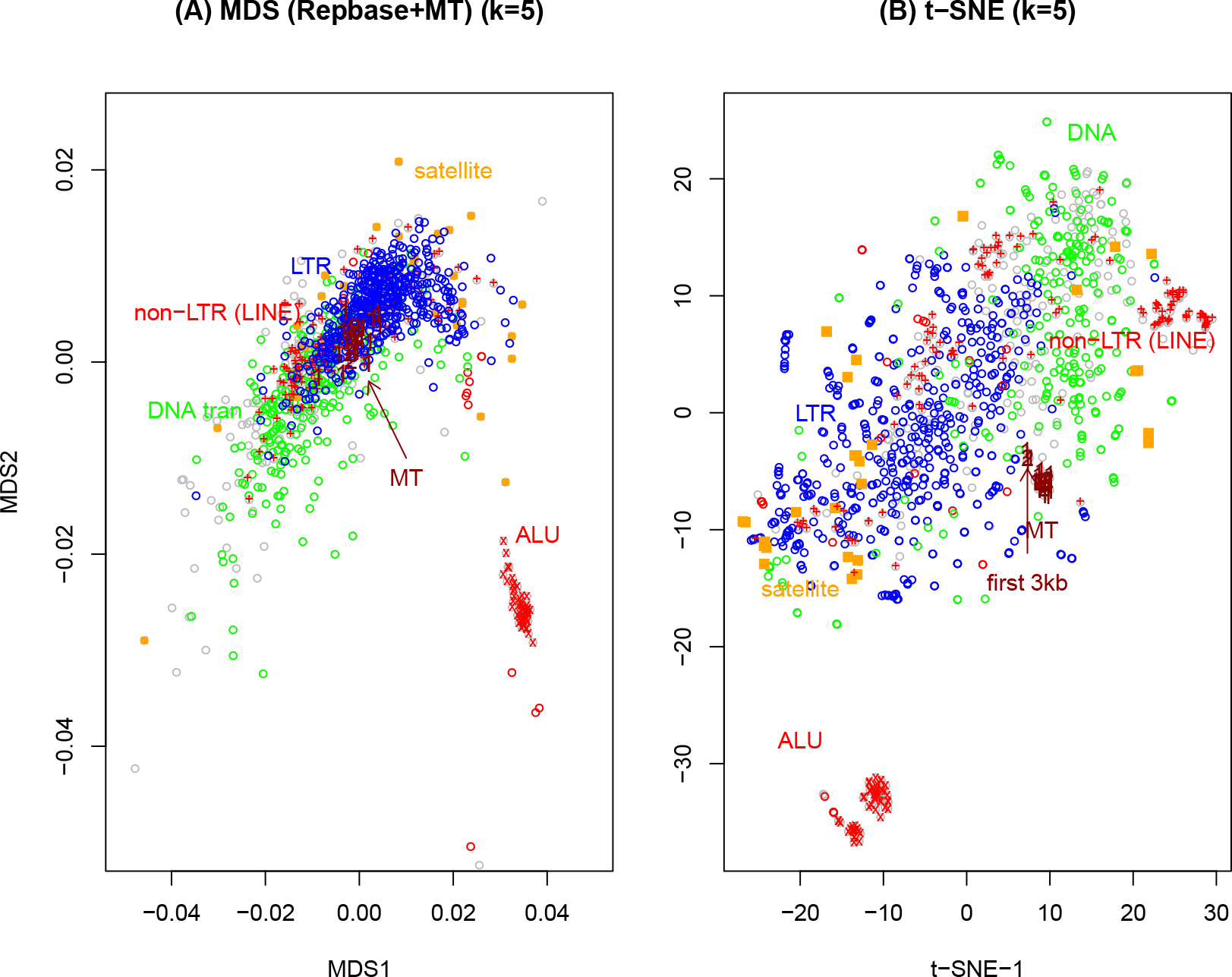
Multi-dimensional scaling (MDS) (A) and t-SNE (B) representation of 5-mer distribution of all sequences in Repbase and various sequences from mitochondria (mitogenome itself, five 3kb sub-sequences, and 16 1kb sub-sequences from MT). The MDS is zoomed in to focus on the main cluster, thus a few outliers (satellite sequences) are not shown. LTR: red, non-LTR: red (the cross × for Alu, plus + for LINE), DNA transposon: green, satellite sequences: orange, MT: dark red. The sequences without specific label are in grey color.

## Discussion

We use a novel alignment-free strategy for predicting NUMTs through sequence similarity. Our search identifies several nuclear genome regions with yet unrecognized sequence similarity to the mitochondrial genome, which may indicate a common ancestry, and not mutually exclusive may indicate potential MT insertions followed by complicated mutational dynamics. The principal validity of our alignment-free approach is supported by the fact that all of our detected NUMTs for the parameters of k=6 and k=7 (and window size 3kb) were also identified with standard alignment-based methods. This confirms a previous conclusion that alignment-free methods can be as powerful as (or even outperform) Smith-Waterman alignment (Zielezinski et al., 2017) for the detection of homologous sequences. The principal question of finding NUMTs is very similar to the situation of detecting repetitive/transposable sequences in the human genome (De Koning et al., 2011), where the result depends on the parameter values chosen, such as proportion of bases in a window that match signature k-mers (k=14-15 in that application). Indeed, our novel similarity signal is only observed when *k* = 5 and *k* = 4.

The motivation of using alignment-free methods in detecting homologous sequences has been well summarized in a recent review (Zielezinski et al., 2017). Our approach mainly aims at the problem (case no.1 in (Zielezinski et al., 2017)) that alignment-based approaches may miss MT insertions, if the insertion event was followed by a more complicated evolutionary dynamics. Although we follow the basic playbook in k-mer distribution approach (Fig.1 in (Zielezinski et al., 2017)), there are some subtle differences: we compare the k-mer distribution from both reference strand and the reverse complement strand of MT with that of the nuclear DNA reference strand, but strand symmetry is not used to combine k-mer frequencies; and we use the JS distance instead of Euclidean distance (because we found that the former leads to a more consistent result with the alignment-based method).

We have chosen our NUMT calling criterion carefully, by using the peak heights from the known (i.e., alignment-based) NUMTs and allowing for a 5% false negative rate, in order to avoid high false positive rates. Although we do not have a theoretical foundation for our choices (which are mostly made by trial-and-error), there are multiple reasons to believe that chance events do not cause observed k-mer (at k=5 or 4) similarity peaks, including those which are not listed among the previously known NUMTs. First of all, when the sequence is reversed (but not complementary), 1/JS values in these peak regions are much reduced. This is another way to say that an insertion followed by an inversion is very rare at the 3kb range or longer. Secondly, when k=3, all peaks observed at k=5 or k=4 spanning more than one window (see Table 2) are remain to exist as peaks. This consistency for reduced k indicates the robustness of the result.

It is not unexpected that using shorter *k*-mers, alignment-free approaches could detect signals that were not seen with alignment-based methods. After all, the purpose of alignment methods is to produce a continuous stretch of sequence which is common in two sequences, whereas alignment-free methods do not concern about the spatial orders of the enriched k-mers. By reducing the seed word length, BLAST may produce more, albeit shorter, alignments, but these shorter alignments are not reported by BLAST as part of a single hit. While no novel sequence similarity peaks are detected for k=6 or k=7, a sub-significance signal also exists for the regions which are significant for k=4 or k=5.

A previous application of alignment-free methods to mitochondrial genome has distinguished different MT haplogroups (with close to 5000 of them existing) (Navarro-Gomez et al., 2014). This is a very different goal from our analysis: all MT haplogroups have similar k-mer distributions when k is low (e.g., k=4-7), so in order to identify one haplogroup among many, the signature k-mers have to be unique to that haplogroup, which can only be achieved with a large k. Accordingly, the choice in (Navarro-Gomez et al., 2014) was to use k=12.

Alignment-free approaches further have the advantage of greater computational speed (Zielezinski et al., 2017), which makes it easy to use our approach to scan NUMTs in the genomes of other species. We easily confirmed the 270kb MT insertion in chromosome 2 of plant Arabidopsis thaliana (plot not shown) (Lin et al., 1999) (another experiment found the insertion size is 620kb (Stupar et al., 2001)). A similar check for chloroplastic DNA showed none of such peaks (data not shown).

To visualize the regions of nuclear DNA that are predicted NUMTs, we use a so-called Manhattan plot. Manhattan plots are standard tools in genome-wide association studies (GWAS) for complex disease, but have been used less in studies that evaluate the evolutionary history and functional content of the human genome. Here we plot the reciprocal of JS over chromosome positions, such that peaks indicate nuclear regions that are similar to the mitochondrial sequences. On the other hand, when JS is plotted instead of 1/JS, the peaks usually indicate centromeres, telomeres, and other low-complexity regions (plot not shown), as these contain distinct repeating patterns (e.g., (Thanos et al., 2018)) Comparing to other graphic illustration of NUMTs (e.g., Fig.2 of (Woischnik and Moraes, 2002)), our approach displays not only the chromosomal location, but also strength of the signal.

The overlap of our novel NUMT predictions with LTR annotations requires some specific discussion. A previous report indicated that flanking region of NUMTs are enriched with retrotransposons (Tsuji et al., 2012). The consistent overlap of our novel predictions with LTR annotations may indicate that alignment-based methods are particular insufficient to identify NUMTs within LTRs, potentially due to certain mutational dynamics contributing to the silencing of repeats. Also the presence of repeats in the mitogenome has been noticed: in the fungal mitogenome, a 17-mer is repeated three times, among other repeats (Misas et al., 2016); and mobile elements were studied in yeast MT (Wu and Hao, 2015); etc. Another potentially interesting observation is that the reverse transcriptase sequences are shown to be similar to fungal introns (Xiong and Eickbush, 1988). Therefore, there is the possibility that certain ancestral transposons were inserted into the mitochondrial genome. Finally there is the possibility that the high degree of sequence similarity does not predict homology, but due to common ancestry. Further careful analyses are need to distinguish various scenarios.

In conclusion, we find that current NUMTs predictions are equally well detected by for alignment-free methods for certain parameter range (e.g. *k* > 5), but additional NUMTs are detected with smaller choices of k (k=5 or k=4). The precise nature of these novel NUMT predictions remains to be determined.

## Data and Methods

### The list of Human NUMT

The list of previously established NUMTs is obtained from the supplementary material of (Calabrese et al., 2012): https://static-content.springer.com/esm/art:10.1186/1471-2105-13-S4-S15/MediaObjects/1285920125112MOESM1ESM.xls. Accord-ing to (Simone et al., 2011; Calabrese et al., 2012), the list was generated by BLASTN 2.2.19 with these score parameters: 2 for match reward, −3 for mismtach, −5 for gap opening, −2 for gap extension, and expected value 1*E* −3. The initial word length is not mentioned in (Simone et al., 2011; Calabrese et al., 2012), but we assume it is k=11 as it is the NCBI BLAST default for blastn. The chromosome positions in this list are in hg19, and we applied the *LiftOver* program (*https://genome.ucsc.edu/cgi-bin/hgLiftOver*) to convert the NUMT coordinates to hg38.

### Code/script

The k-mer count programs are implemented as custom scripts in Python and Perl by the authors. The plot and analysis was carried out in R (*https://www.r-project.org/*), including the classic multi-dimensional scaling cmdscale() and t-distributed stochastic neighboring embedding Rtsne() (*https://github.com/jkrijthe/Rtsne*).

### Alignment methods

We used (i) the BLAST web interface at NCBI *https://blast.ncbi.nlm.nih.gov/Blast.cgi*, (ii) command line run by a local BLAST+ 2.7.1 copy (*ftp://ftp.ncbi.nlm.nih.gov/blast/executables/blast+/LATEST/*), and (iii) the LAST (Kielbasa et al., 2011) alignment program, (*http://last.cbrc.jp/*) to obtain pairwise alignments.

### BLAST

For the NCBI BLAST web interface, we choose “Nucleotide BLAST” → “Align two or more sequences” → (parameter setting) “Somewhat similar sequences” (blastn). This would choose the default parameter settings: match/mismatch/gap-open/gap-extension score same as in (Simone et al., 2011; Calabrese et al., 2012), expected value threshold of 10, and k=11. We also varied k from 11 to 7 (there are only three choices: 7, 11, and 15), and other higher expected threshold values.

For the local copy of blastn as part of the BLAST+ package, the default score for match/mismatch/gap-open/gap-extension is 1, −2, 0, and −2.5. The initial seed word length is changed by the −word size command option, but it has to larger than 4 (default value is 11). The expected value is changed by the −evalue option (default value is 10).

### LAST

The LAST alignment program (Kielbasa et al., 2011) does not require the tuning of the seed word length *k*, because it starts from single nucleotides (*k* = 1). The threshold can be adjusted by several parameters, and one is *D* (default value for *D* is 20000). LAST report alignments that are expected by chance at most once per D-length of the query sequence. The default match/mismatch/gap-open/gap-extension score in LAST is: 2, −3, −7, −1.

### Genome annotation

We used the UCSC Genome Browser *https://genome.ucsc.edu* to obtain annotation at queried chromosomal regions, choosing “(Genomes) Human GRCh38/hg38”. Both mitochondrial and nuclear DNA sequences were downloaded from *http://hgdownload.soe.ucsc.edu/goldenPath/hg38/chromosomes/* (chr1.fa.gz, chr2.fa.gz, · · · chrM.fa.gz).

### Repbase database

the database of repetitive DNA elements, Repbase (Jurka, 2000; Kojima, 2018), is downloaded from *https://www.girinst.org/repbase/* in November 2016. We use the two FASTA sequence files: humrep.ref (1043 sequences) and humsub.ref (71 sequences).

## Appendix

### BLAST and LAST alignment between new alignment-free peak regions at chromosomes 1,3 and the mitogenome

BLAST(online) has these default parameter values: scores for match/mismatch/gap-open/gap-extension are 2, −3, −5, −2, seed word length *k* = 11, and expected threshold *E* = 10. There are only three options for *k* = 7, 11, 15. BLAST(local) has scores for match/mismatch/gap-open/gap-extension 1, −2, 0, −2.5, and *k* ≥ 4 can be adjusted, as well as *E*. The default setting for match/mismatch/gap-open/gap-extension score in LAST is 2, −3, −7, −1, and default threshold is *D* = 20000.

The following two tables illustrate the BLAST(online), BLAST(local), LAST alignment results between chr1:9087750-9096750 (hg38) or chr3:141694900-141702000 and human mitochondrial DNA, at the default score parameter values and some of the allowed *k* and *E* values. The results include the number of alignments that pass the threshold (#match), the longest alignments (maxL), sum of scores over all alignments (totalS).

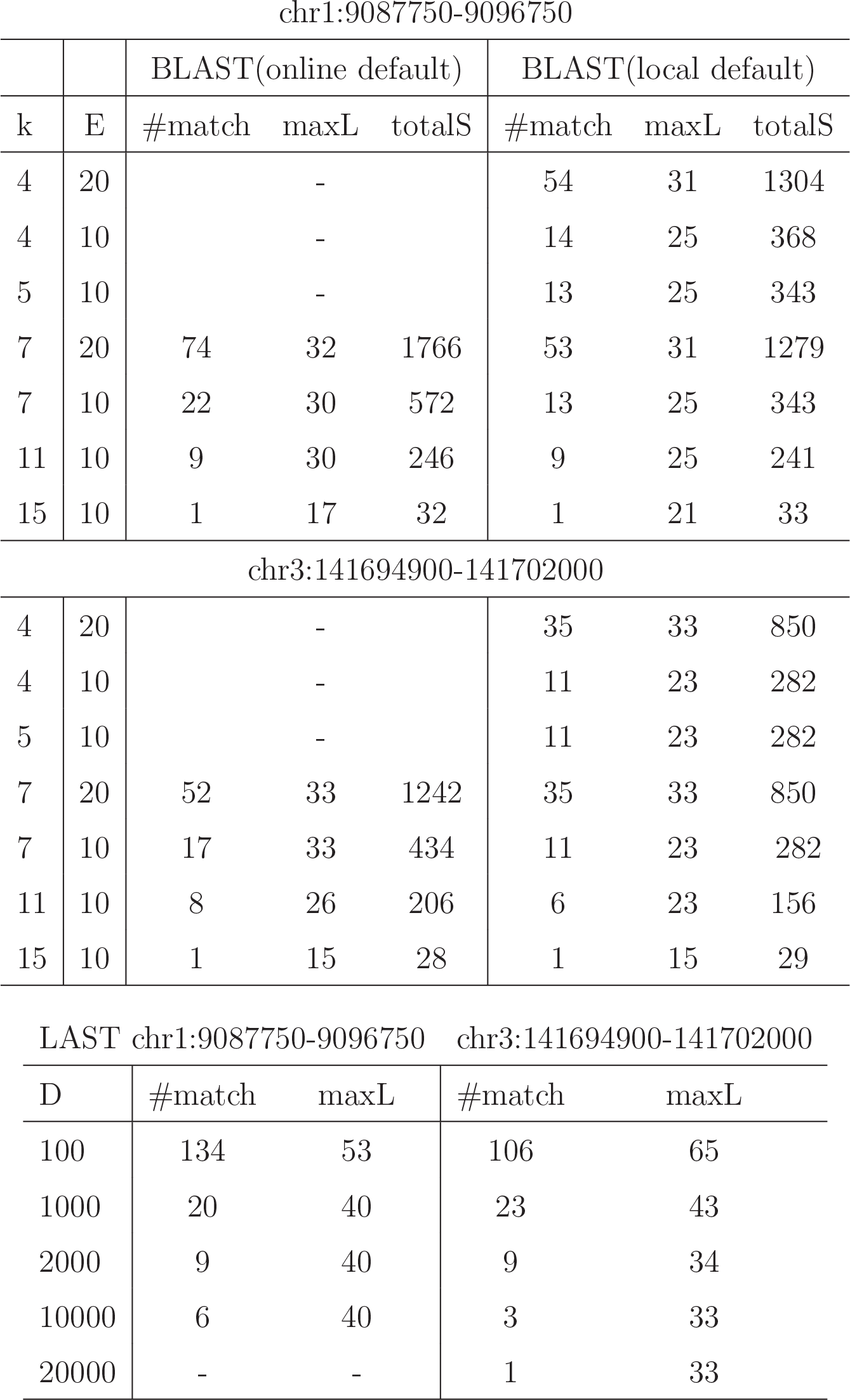

|  |  | BLAST(online default) |  |  | BLAST(local default) |  |  |
| --- | --- | --- | --- | --- | --- | --- | --- |
| k | E | #match | maxL | totalS | #match | maxL | totalS |
| 4 | 20 |  | - |  | 54 | 31 | 1304 |
| 4 | 10 |  | - |  | 14 | 25 | 368 |
| 5 | 10 |  | - |  | 13 | 25 | 343 |
| 7 | 20 | 74 | 32 | 1766 | 53 | 31 | 1279 |
| 7 | 10 | 22 | 30 | 572 | 13 | 25 | 343 |
| 11 | 10 | 9 | 30 | 246 | 9 | 25 | 241 |
| 15 | 10 | 1 | 17 | 32 | 1 | 21 | 33 |
| chr3:141694900-141702000 |  |  |  |  |  |  |  |
| 4 | 20 |  | - |  | 35 | 33 | 850 |
| 4 | 10 |  | - |  | 11 | 23 | 282 |
| 5 | 10 |  | - |  | 11 | 23 | 282 |
| 7 | 20 | 52 | 33 | 1242 | 35 | 33 | 850 |
| 7 | 10 | 17 | 33 | 434 | 11 | 23 | 282 |
| 11 | 10 | 8 | 26 | 206 | 6 | 23 | 156 |
| 15 | 10 | 1 | 15 | 28 | 1 | 15 | 29 |

LAST chr1:9087750-9096750 chr3:141694900-141702000
| D | #match | maxL | #match | maxL |
| --- | --- | --- | --- | --- |
| 100 | 134 | 53 | 106 | 65 |
| 1000 | 20 | 40 | 23 | 43 |
| 2000 | 9 | 40 | 9 | 34 |
| 10000 | 6 | 40 | 3 | 33 |
| 20000 | - | - | 1 | 33 |

## Acknowledgment

Jerome F would like to thank the financial and other support from the Summer Internship Program of the Feinstein Institute for Medical Research. We thank Andrew Shih, Susana Marquez Renteria, Ilya Korsunsky, Jane Cerise, Daniel Miller, Oliver Clay, Dimitris Thanos, Astero Provata, and Pedro Miramontes for discussions.

